# A versatile resource of 1500 diverse wild and cultivated soybean genomes for post-genomics research

**DOI:** 10.1101/2020.11.16.383950

**Authors:** Hengyou Zhang, He Jiang, Zhenbin Hu, Qijian Song, Yong-qiang Charles An

## Abstract

With the advance of next-generation sequencing technologies, over 15 terabytes of raw soybean genome sequencing data were generated and made available in the public. To develop a consolidated, diverse, and user-friendly genomic resource to facilitate post-genomic research, we sequenced 91 highly diverse wild soybean genomes representing the entire US collection of wild soybean accessions to increase the genetic diversity of the sequenced genomes. Having integrated and analyzed the sequencing data with the public data, we identified and annotated 32 million single nucleotide polymorphisms (32mSNPs) with a resolution of 30 SNPs/kb and 12 non-synonymous SNPs/gene in 1,556 accessions (1.5K). Population structure analysis showed that the 1.5K accessions represent the genetic diversity of the 20,087 (20K) soybean accessions in the U.S. collection. Inclusion of wild soybean genomes significantly increased the genetic diversity and shorten linkage disequilibrium distance in the panel of soybean accessions. We identified a collection of paired accessions sharing the highest genomic identity between the 1.5K and 20K accessions as genomically “equivalent” accessions to maximize the use of the genome sequences. We demonstrated that the 32mSNPs in the 1.5K accessions can be effectively used for in-silico genotyping, discovering trait QTL, gene alleles/mutations, identifying germplasms containing beneficial allele and domestication selection of trait alleles. We made the 32mSNPs and 1.5K accessions with detailed annotation available at SoyBase and Ag Data Commons. The dataset could serve as a versatile resource to release the potential of the huge amount of genome sequencing data for a variety of postgenomic research.

## Introduction

Soybean [*Glycine max* (L.) Merr.] is one of the most economically important field crops for its high seed protein (40%) and oil (20%), which are primarily used for animal feed, human consumption, and industrial use. Soybean can also fix atmospheric nitrogen and plays an important and sustainable role in agriculture. Although worldwide soybean production has been tripled and its growing acreage has doubled since 1993 (USDA-OCE 2017), it is predicted that demand for soybean continuously increases for a growing world population and for better nutritional value (Tilman *et al*., 2011).

The first complete soybean genome was available in 2010 (Schmutz *et al*., 2010). Since then, with the continuous reduction in sequencing cost, whole genomes of many soybean accessions have been re-sequenced (Kim *et al*., 2010; Lam *et al*., 2010b; Valliyodan *et al*., 2016; Zhou *et al*., 2015; Fang *et al*., 2017). So far, more than 1,400 whole soybean genomes have been re-sequenced and are available in a format of raw sequencing reads in the public databases, such as Sequence Read Archive (SRA) of the NCBI (https://www.ncbi.nlm.nih.gov/sra). Most of those soybean accessions are important for soybean basic and applied research such as elite cultivars, parents of genetic mapping populations, and germplasm containing beneficial alleles for soybean improvement. Availability of the sequencing data provides an unprecedented opportunity to access the genome sequences of those individual germplasms or explore their soybean genomic diversity in a large population (Zhou *et al*. 2015; Li *et al*., 2014). Unfortunately, it remains a big challenge for most laboratories to systemically analyze such a huge amount of raw genome sequencing reads, thus these valuable re-sequencing data are under-utilized. There is a pressing need to analyze those genome sequencing data and generate a public-accessible resource in a user-friendly format to release the huge potential of those genome sequences for their research.

Soybean (*G. max*) was domesticated from *G. soja* in China about 5,000 years ago (Hymowitz 1970). However, domestication and modern breeding dramatically reduce the genetic diversity in modern soybean cultivars. For example, current soybean cultivars in North America have lost 28% of the sequence diversity in the Asian landraces and 81% of the rare alleles present in the wild soybean (Hyten *et al*., 2006). Loss of the critical genetic diversity and beneficial alleles in cultivated soybean population imposes a challenge for continuous soybean improvement with new trait genes/variation. In addition, those aforementioned projects of soybean genome sequencing mainly focus on cultivated soybean, with a limited number of wild soybean accessions being sequenced and analyzed. There is a strong need to sequence additional highly diverse wild soybean accessions to enrich gene pool of the sequenced genomes for discovering novel and superior trait genes and alleles (Kofsky *et al*., 2018; Stupar 2010; Qi *et al*., 2014).

In the study, we sequenced 91 *G. soja* accessions representing the diversity of *G. soja* in the US Soybean Germplasm Collection and integrated them with 1,465 publicly available re-sequenced genome reads. Our analysis identified and annotated 32 million SNPs (32mSNPs) present in a total of 1,556 (1.5K) diverse soybean germplasm (*G. soja*, landrace, improved cultivar) collected worldwide. In addition, we characterized and provided a comprehensive description of the population structures and biological diversity of the 1.5K accessions. We showed that the collection of 1.5K accessions represents the genetic diversity of over 20,087 (20K) soybean accessions in the current US Soybean Germplasm Collection. The collection of 32mSNPs in 1.5K accessions should serve as a valuable resource for using the untapped value of the huge amount of soybean genome sequencing data and exploring the genome diversity of the 1.5K diverse soybean accessions for various soybean basic and applied research.

## Results

### A collection of 1,556 diverse soybean accessions

We sequenced whole genomes of 91 diverse *G. soja* accessions at a depth ranging from 7.4 to 41.3-fold genome coverage. The 91 *G. soja* accessions represent the overall diversity of 1,168 *G. soja* accessions in the US Soybean Germplasm Collection based on their genetic distances, geographic locations, and maturity groups (Song *et al*., 2015a). In addition, we retrieved whole-genome sequencing reads of 1,465 genomes from the NCBI SRA database with their sequencing depth ranging from 2.7-to 65.1-fold genome coverage (**Table S1**). The genome sequencing reads of 1,556 (hereafter referred to as 1.5K) diverse soybean genomes were combined for the following analysis.

This collection consisted of 204 *G. soja* and 1,352 *G. max* accessions. Out of the 1.5K accessions, a total of 1,194 accessions were annotated for their germplasm types, including 204 *G. soja* accessions, 472 landraces, and 518 improved cultivars (**Table S1**). The 1.5K accessions were collected widely across the world including major soybean growing countries (**Figure 1a and Table 1**). Accessions from China and the United States accounted for the majority of the accessions (71.3%) with the most from China (955 accessions) followed by the United States (155 accessions). The remaining 28.7% (446 accessions) accessions were collected from the other 37 countries. This collection also included 32 accessions from Brazil, thus far the largest

**Figure 1.**
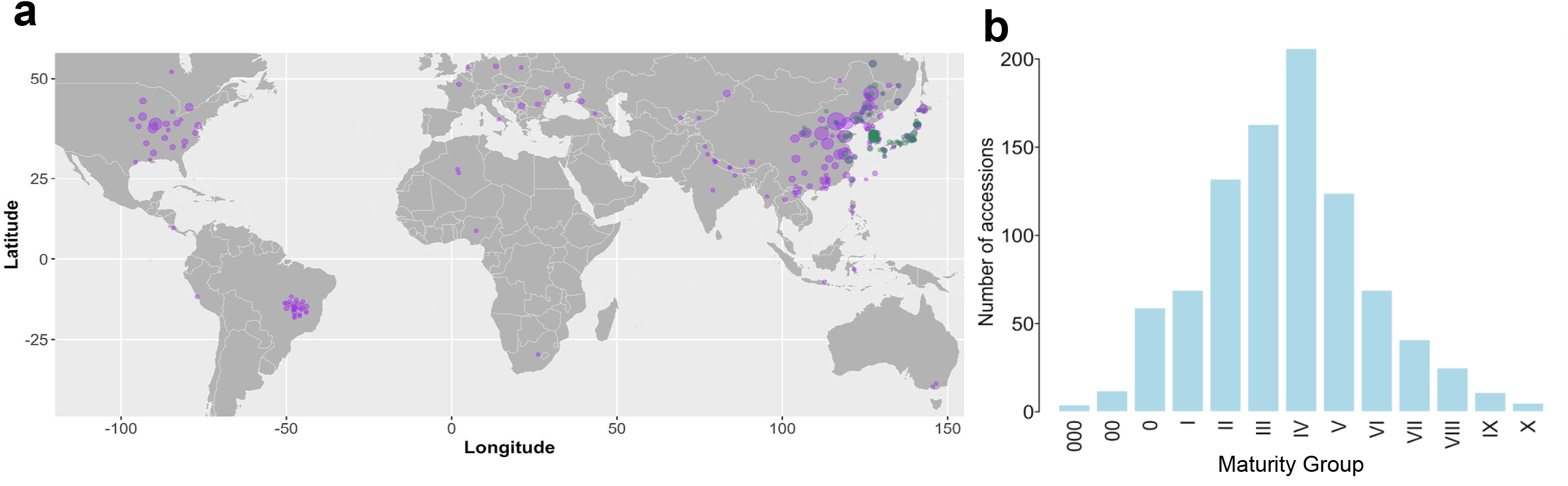
Origins and maturity groups of the 1.5K soybean accessions. **a)** Geographic distribution of the 1.5K collection worldwide. The dots in green denote *G. soja* and those in purple denote *G. max*. Dot size denotes the sample size at the indicated geographic region. **b)** Distribution of maturity group for all the accessions. Each bar represents the number of accessions belonging to the corresponding maturity group.

**Table 1.** The origins of 1.5K soybean collection

| Origins | No. of<br>accessions | Origins | No. of<br>accessions |
| --- | --- | --- | --- |
| China | 955 | Nepal | 2 |
| United States | 155 | Former Serbia and<br>Montenegro | 2 |
| Japan | 105 | Serbia | 2 |
| South Korea | 93 | Indonesia | 2 |
| Russian Federation | 42 | Netherlands | 1 |
| Brazil | 32 | Peru | 1 |
| North Korea | 17 | Georgia | 1 |
| Canada | 14 | Kyrgyzstan | 1 |
| India | 8 | Belgium | 1 |
| Vietnam | 6 | Italy | 1 |
| Moldova | 4 | Nigeria | 1 |
| Ukraine | 3 | Myanmar | 1 |
| Germany | 3 | South Africa | 1 |
| Romania | 3 | Uzbekistan | 1 |
| Algeria | 2 | Sweden | 1 |
| France | 2 | Costa Rica | 1 |
| Philippines | 2 | Austria | 1 |
| Australia | 2 | Poland | 1 |
| Thailand | 2 | Unknown | 27 |
| Hungary | 2 | Total | 1501 |

Note: Duplicated-sequenced genomes were counted once here.

soybean producer (USDA-FAS 2020). All the *G. soja* accessions originated from East Asian countries (Russia, China, South Korea, Japan). In addition, the 1.5K accessions were adaptive to all of the thirteen maturity groups (MGs 000-X) and covered all soybean cultivation areas with the majority (67.93%) of the collection in MGs II, III, IV, and V (**Figure 1b** and **Table S1**). Thus, this 1.5K collection comprised diverse accessions worldwide that should contain abundant genetic diversity in soybean, especially the untapped diversity retained in wild soybean.

### Identification and annotation of 32mSNPs in the 1.5K diverse soybean accessions

We analyzed a total of 208.3 billion paired-end and single-end sequence reads of the 1,556 genomes with an average of 133 million reads and 14-fold genome coverage per accession (**Table S1**). A total of 32,456,244 SNPs (designate 32mSNPs) were identified at an average SNP density of 30 SNPs/kb. 16.3 million and 8.4 million SNPs had minor allele frequency (MAF) higher than 0.01 and 0.05 respectively. We observed a higher density of SNPs in euchromatic regions than in heterochromatic regions (**Figure 2a**). Approximately, 87% of the 32mSNPs were located in intergenic regions. The rest 13% (5,193,083) were in the genic regions with an average density of 93 SNPs per gene (**Figure 2b**). Of the genic SNPs, 12.1% and 63.1% were located in untranslated regions (5’ and 3’UTR) and introns respectively (**Figure 2c**). The remaining 24.8% of the genic SNPs were present in the coding sequences (CDS) (**Figure 2c**). We observed that 63% (657,371) of these SNPs in the CDS were non-synonymous. PROVEN algorithm predicted 10.7% (70,143) of the non-synonymous SNPs as deleterious with PROVEN score ≤ 4.1, which have a high possibility in altering gene functions (Choi *et al*., 2012). In addition, we identified 19,506 SNPs at splicing sites, 1,562 SNPs at start codons, 1,465 SNPs at stop codons, and 22,076 SNPs producing premature stop codons (**Figure 2d**). Importantly, 99% of the 56,044 gene models in the soybean genome (*Wm82*.*a2 v1*) carried at least one non-synonymous SNP with an average of 12 non-synonymous SNPs/per gene. Thus, the collection of 1.5K accessions are highly diverse, and likely contained at least one mutation/allele almost for each gene that potentially alter gene functions and cause phenotypic variation, which allows to explore those allelic variants and associated phenotypic variations in 1.5K soybean accessions to discover novel genes and alleles important for soybean improvement.

**Figure 2.**
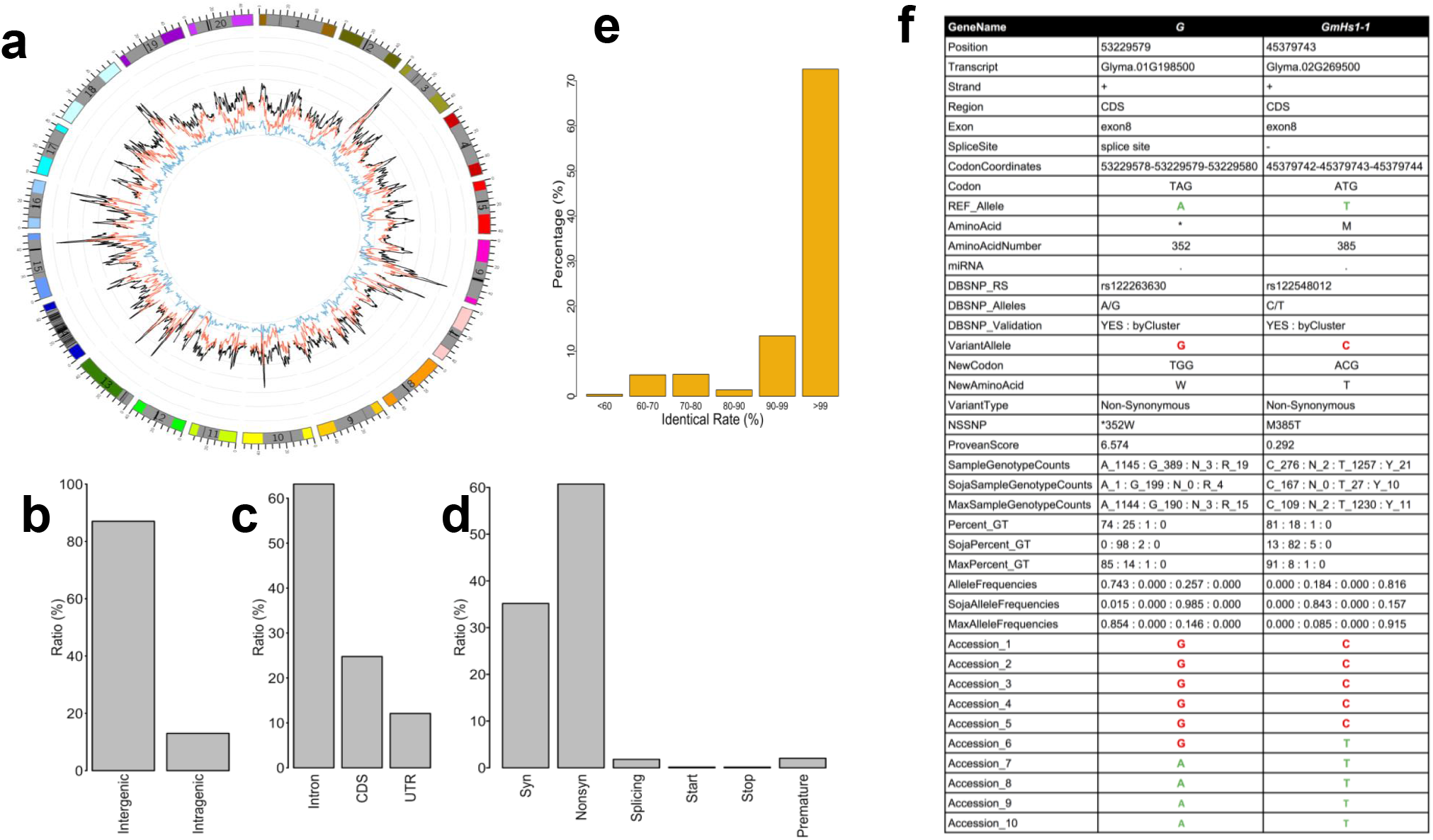
Annotation of the 32mSNPs. **a)** Distribution of the SNPs along 20 chromosomes compared with the dbSNPs. The outermost circle represents the 20 soybean chromosomes. 0, 20, and 40 outside the circle represent 0 Mb, 20 Mb, and 40 Mb positions on the chromosome, respectively. The solid gray boxes and black bars indicate pericentromeric regions and centromeric repeats, respectively. The black, red, and blue curves of the inner circle showed the distribution of the SNP set identified in the 1.5K accessions, soybean dbSNPs, and the subset of SNPs with minor allele frequency ≥ 0.25, respectively. **b)** Percent of the SNPs in the 32mSNPs in genome features (intergenic and intragenic). **c)** Percent of genic SNPs in the 32mSNPs (Intron, CDS, UTR). **d**) Percent of synonymous (Syn), non-synonymous (Nonsyn) SNPs, and the SNPs in splicing sites, start codons, stop codons, and the SNPs causing premature in the 32mSNPs. **e**) Percent of identity at the coordinate-matched SNPs for the accessions that have been genotyped by genome-resequencing and SoySNP50K. **f**) A SNP report exemplified with two previously identified domestication genes. *G* encodes a stay-green gene controlling soybean dormancy and *GmHs1-1* encodes calcineurin-like protein controlling har-seededness. The SNP report contains 30 annotation categories for each identified SNP in the 32mSNPs. Reference and alternative alleles were highlighted in green and red, respectively.

Fifty-six accessions have been sequenced twice in different studies. The duplicated sequences for the same accessions had an average of 99.5% of sequence identity, indicating high reproducibility and high accuracy of the SNP calling in the present study. Out of the 1.5K accessions, 926 accessions were also genotyped using Soy50KSNP Chip in a separate study (Song *et al*., 2015b) and about 72.57% of 926 accessions had > 99% of identity at SNP positions genotyped by both platforms (**Figure 2e**). The remaining 27% of the accessions showed less than 99% of identity between the two platforms. Having compared the 32mSNPs with 15,623,492 registered soybean SNPs available in the NCBI dbSNP database (https://www.ncbi.nlm.nih.gov/snp/), we observed that 13 million SNPs were present in the dbSNP database. Thus, 19 million out of the 32mSNPs represented novel SNPs.

To provide a comprehensive description of each SNP, especially those non-synonymous SNPs potentially important to gene functions in the 1.5K collection, we structurally and functionally annotated each SNP with 30 categories of information such as the SNP position, reference and alternative alleles, and their allelic frequency in *G. soja* and *G. max*. (**Figure 2f**). *G* and *GmHs1-1*, are two agronomically important genes (Wang *et al*., 2018; Sun *et al*., 2015) (**Figure 2f**). *GmHs1-1* encodes a calcineurin-like protein controlling hard-seededness and *G* gene controls soybean dormancy in soybean. The annotation showed that the causative reference and alternative alleles for *GmHs1-1* (reference T/ alternative C) and *G* (A/G) were precisely identified in the 1.5K collection with an average of 98.6% of the 1.5K carrying homozygote calls for the alleles. The annotated percentages of each allele in subpopulations (*G. max* and *G. soja*) showed that highly biased presences of the two alleles in *G. soja* and *G. max* for each of the two genes. It is consistent with the previous reports that both were domestication genes. For example, the *G* gene had 85% of A in *G. max* and 98% of G in *G. soja*. Thus, the comprehensive annotation is highly valuable for the post-genomic research, such as population-scale characterization of genome-wide SNPs and genes for discovering domestication-associated SNPs or genomic regions of interest. These results further exemplified the high coverage and robustness of the 32mSNPs and its application in characterizing genes and its association with traits.

### Population structure of 1.5K soybean accessions

We assessed the population structure using principal component analysis (PCA) and neighbor-joining (NJ) phylogenetic analysis. The PCA analysis revealed that the first 20 principal components (PCs) of the genetic data captured 32.53% of the total variance in the 1.5K accessions. The first PC captured 9.25% of the variation, mainly explaining the divergence between *G. max* and *G. soja* (**Figure 3a**). The second PC captured 5.23% of the variation, mainly explaining variation within the germplasms, and the third PC captured 3.09% variation and mainly explained the variation within the remaining variation within the germplasms (**Figure 3b**). We further constructed a neighbor-joining phylogenetic tree using the 1.5K accessions. The tree showed consistent results as observed in PCA, in which accessions were clustered based on the germplasm types with some admixture between germplasms (**Figure 3c**). Especially, landraces and cultivars preferentially clustered together respectively but no clear pattern was observed between the clusters, suggesting the existence of genetic diversity shared between these landrace and cultivar accessions. However, *G. soja* accessions were mainly clustered exterior to *G. max* clusters. Therefore, this collection has a clear, germplasm type-inferred population structure that allows to leverage the high-resolution SNPs to investigate complex processes of soybean domestication and improvement.

**Figure 3.**
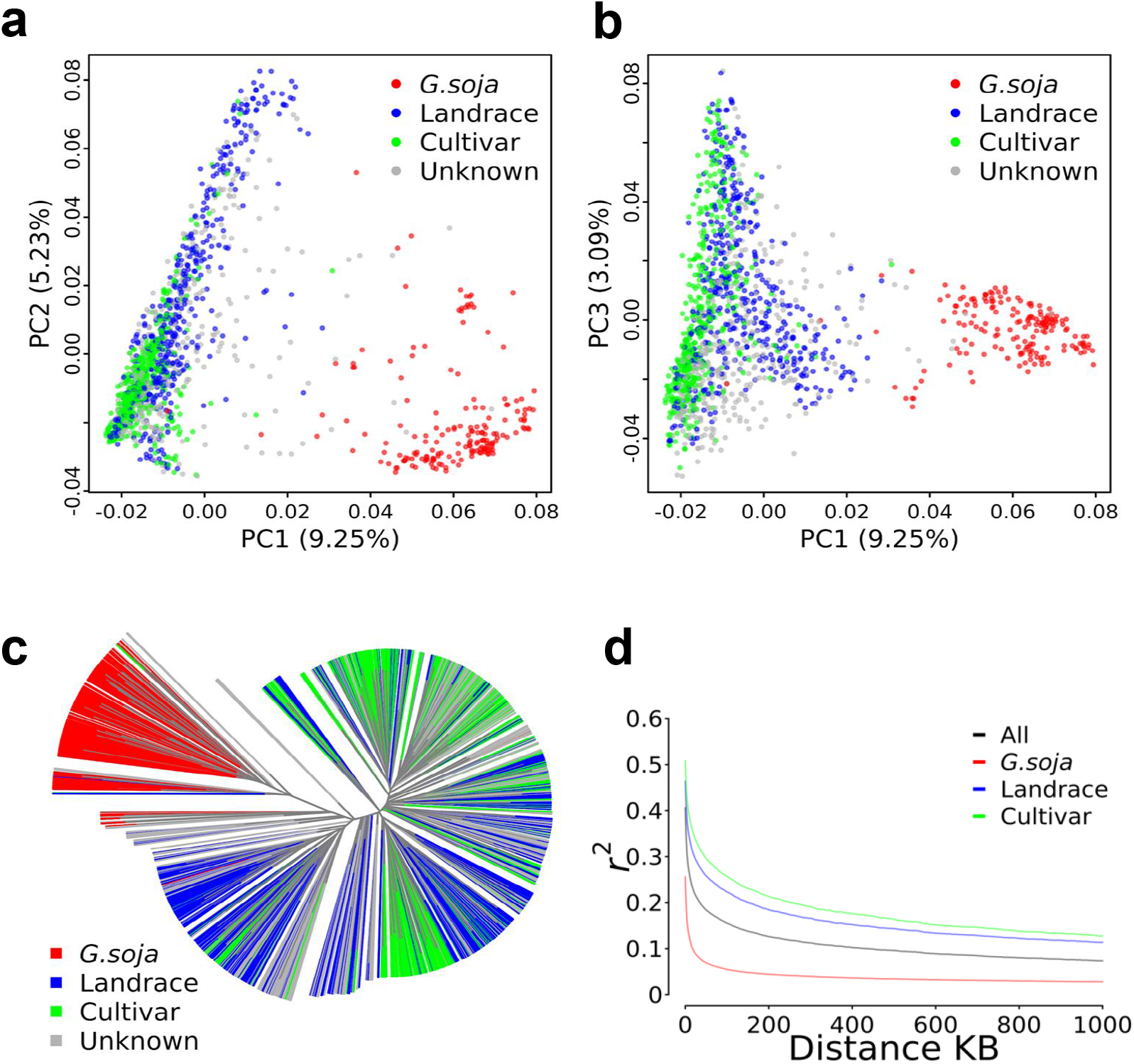
Population genomic analysis of the 1.5K collection. **a)** Scatter plots for the first two components (PC1, PC2) of the 1,556 (1.5K) accessions, and **b**) for PC1 and PC3. **c**) A neighbor-joining phylogenetic tree of the 1.5K accessions. The germplasm types (*G. soja*, landrace, cultivar) are correspondingly labeled. **d**) Comparison of LD decay among three germplasm types (*G. soja*, landrace, cultivar).

Having estimated linkage disequilibrium (LD) across the three germplasm types (**Figure 3d)**, we observed an overall rapid LD decay in *G. soja* compared to domesticated soybean (landrace and cultivar). LD decay for all accessions (all three germplasm types) was ∼35 KB at *r*^*2*^ of 0.2. The LD decay distance in *G. soja* was ∼2 kb at *r*^*2*^ of 0.2, which was dramatically shorter than ∼151 kb for landraces and much shorter than ∼255kb for cultivars (**Figure 3d**). This result indicates that including highly diverse *G. soja* accessions into the collection greatly reduced the sizes of genome-wide LD blocks, thereby, it should significantly increase the resolution of association mapping by breaking long LD in *G. max* population alone.

### The 1.5K collection represents genetic diversity of 20K US Soybean Collection

To determine if high genetic diversity in the 1.5K diverse collection can represent those in the 20K accessions in the US Soybean Germplasm Collection (wild and cultivated accessions) that have been genotyped by SoySNP50K Chip, we assessed the genetic distribution of the 1.5K accessions among 20K accessions. The PCA analysis showed that the 1.5K accessions evenly spread in major clusters among the 20K accessions (**Figure 4a**). This observation was supported by the presence of the 1.5K accessions (orange) in almost all different clusters in the neighbor-joining phylogenetic tree with the 20K accessions as tree background (gray) (**Figure 4b**). Thus, the 1.5K collection contained the genetic diversity representative to those 20K accessions in the US Soybean Collection, and the 32mSNPs likely represented the most SNPs present in the Collection. Having conducted a pairwise comparison between the two collections of accessions, we identified a genetic “equivalent” accession in the 1.5K accessions that shared the highest sequence identity for each of the 20K accessions (**Figure 4c, Table S1**). The results allow to identify an “equivalent” accession in US collection for an inaccessible sequenced accession or infer the genome sequence of the un-sequenced accessions for maximizing the use of the genome sequences (**Table S2**).

**Figure 4.**
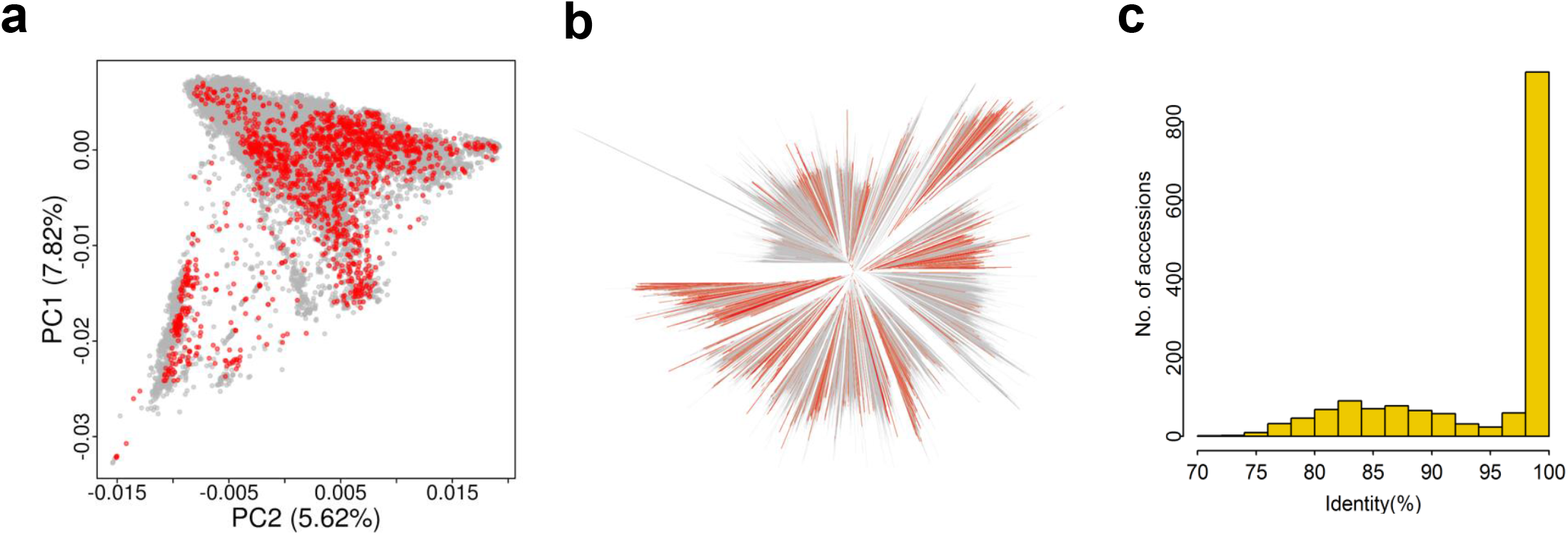
Representation of the 1,556 accessions in the US 20K Soybean Collection. **a)** Distribution of the whole-genome sequenced 1,556 (1.5K) accessions and the 20,087 (20K) accessions using principal component analysis. Red dots denote the 1.5K accessions and gray dots denote the 20K accessions. **b)** A neighbor-joining phylogenetic tree constructed by the 1.5K accessions and 20K accessions. Branches in orange denote the 1.5K whole genomes sequenced accessions and those in gray denote the 20K accessions. **c)** The genomic similarity between the 1.5K accessions corresponding to the 20K accessions as determined by comparing the identity of coordinated-matched SNP sites.

### Application of the 32mSNPs from 1.5K soybean genomes for diverse usages

The 32mSNPs in 1.5K diverse wild and domesticated soybean offers an excellent resource for a variety of post-genomic research. Here we demonstrated that it can be effectively used for forward and reverse genetic research.

#### Forward genetics to fine map trait QTLs

SNPs have served as one of the most widely used molecular markers in genetic research. The 32mSNPs of the 1.5 diverse soybean accessions enables us to leverage such high-resolution SNPs in such a large and diverse population to effectively locate QTL for traits of interest using genome-wide association study (GWAS). Determinate growth habit was phenotyped in 642 accessions out of the 1.5K accessions. GWAS with the 642 accessions revealed two major QTLs with strong associations (*p* < 7.10 × e^8^) on chromosomes 3 and 19 (**Figure 5a**). At the QTL on chr19, a cluster of SNPs spanning 200kb (chr19: 45105190-45305190) was significantly associated with determinate growth. The significant associated SNPs were co-located with a previously reported *Dt1* gene (*Glyma*.*19G194300*) underlying a stem determination QTL (Tian *et al*., 2010; Lam *et al*., 2010a) (**Figure 5c**). Interestingly, a cluster of strongly associated SNPs at the QTL region on chr 3 nearly coincided with a *Dt1* paralog (*Glyma*.*03G194700*) (**Figure 5b**). It had been proposed to be subfunctionalized or neofunctionalized (Tian *et al*. 2010). The GWAS result strongly supports that the *Dt1* paralog likely remains its function in regulating the stem determination after gene duplication. Successfully pinpointing the determinate growth QTLs to the previously identified *Dt1* gene and its paralog with 32mSNPs-inferred GWAS demonstrates the high effectiveness of the large SNP dataset for fine association analysis of a trait of interest to discover their causative QTL genes.

**Figure 5.**
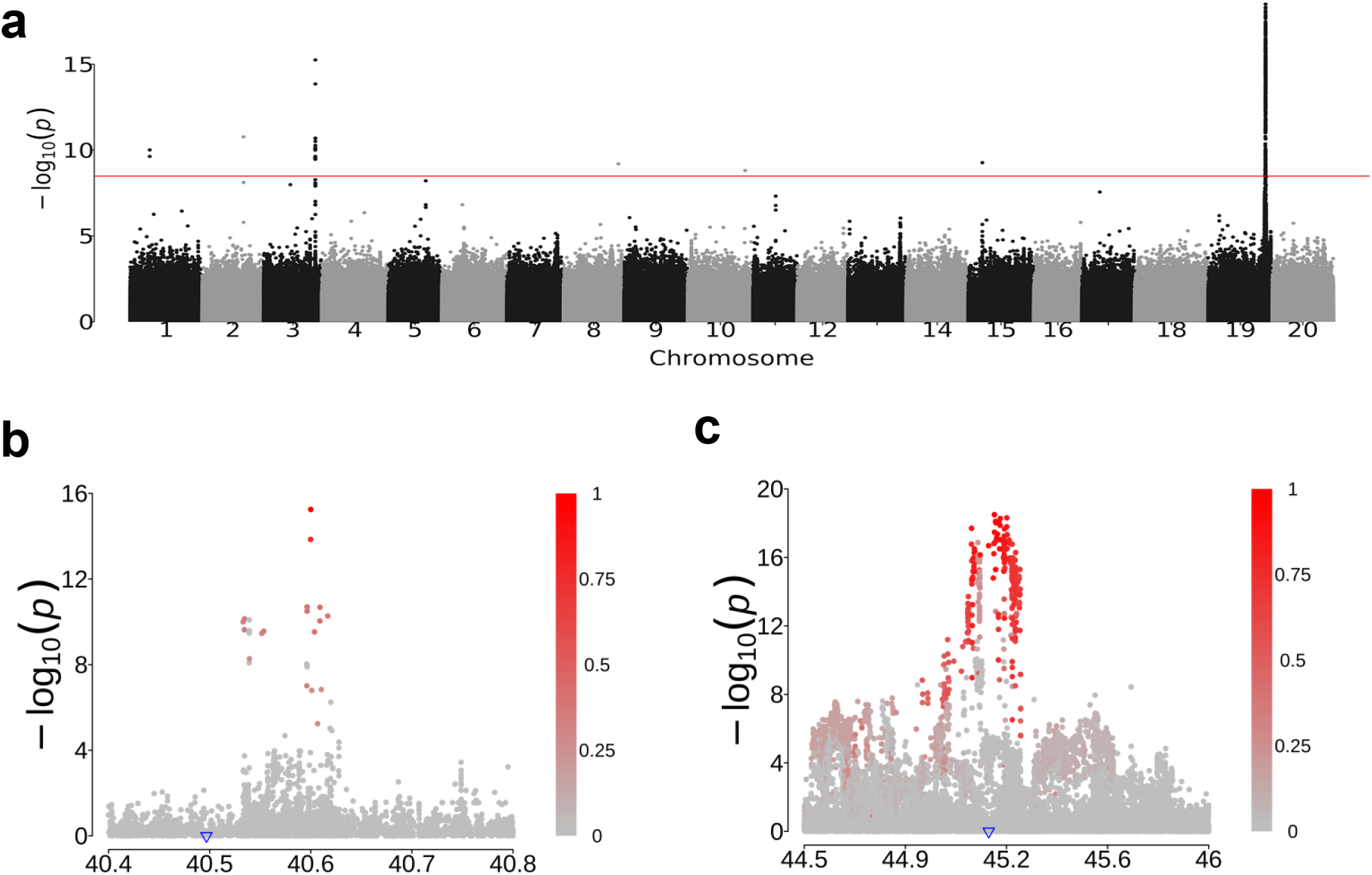
Two associations for stem determinate using the 32mSNPs. **a**) A Manhattan plot illustrating two major QTLs associated with stem determinate on chromosomes 3 and 19. **b**) and **c**) Zoomed-in Manhattan plot for the association regions on chr3 and chr19, respectively. Blue triangles denote the physical positions of the *Dt1* and its homologous gene. The color intensity of each SNP indicates its *r*^2^ value with the peak association SNP. The color scale was shown beside the panel.

#### Reverse genetics to explore the diversity of trait genes

Nucleotide substitution is one of the major DNA variants leading to gene functional and plant phenotypic variation. The 32mSNPs in 1.5K diverse soybeans with comprehensive structural and functional annotation offers an excellent resource for reverse genetic study to identify trait gene mutants present in the 1.5K accessions and germplasm carrying beneficial alleles/variants. It allows us to apply a facile *in-silico* approach with little cost to genotype the SNPs in given genes or defined genomic regions for traits of interest in 1.5K soybean accessions. For example, *FATTY ACID DESATURASE 2* (FAD2) is a key enzyme in fatty acid biosynthetic pathways and plays an important role in regulating fatty acid profile in soybean seeds. In soybean, FAD2 enzymes are encoded by seven *FAD2* genes (Lakhssassi *et al*., 2017). We identified a total of 340 SNPs in *FAD2* genes. Out of the 340 SNPs, 53 non-synonymous SNPs, 2 nonsense SNPs, and 2 SNPs at splicing sites were identified (**Table 2**). These SNPs potentially alter the activities of the genes and seed fatty acid profiles, and likely cause high diversity of seed fatty acid profile in the soybean population. Four SNPs (S86F, M126V, P137R, and I143T) were previously demonstrated to be highly correlated with high oleic acid content (Pham *et al*., 2010; Schlueter *et al*., 2007). We identified three (S86F, M126V, P137R) of the four SNPs, which also reflects a high diversity and coverage of 1.5K collection. Of these variants, the missense mutation P137R in *FAD2-1B* of PI283327 (Pingtung Pearl, maturity group V) has served as a critical allele to breed high oleic acid (Pham *et al*. 2010). Importantly, despite rare alleles, we also successfully identified another accession (PI506933, Kouiku at maturity group IV) carrying the same P137R allele (**Table 3**). This result suggested the importance of preserving all identified variation including rare alleles in the SNP collection. These analyses demonstrated that a combination of the highly diverse soybean accessions with the *in-silico* genotyping analysis offers a highly effective approach to discover new trait alleles and germplasm containing previously identified or new trait gene mutations/alleles.

**Table 2.**
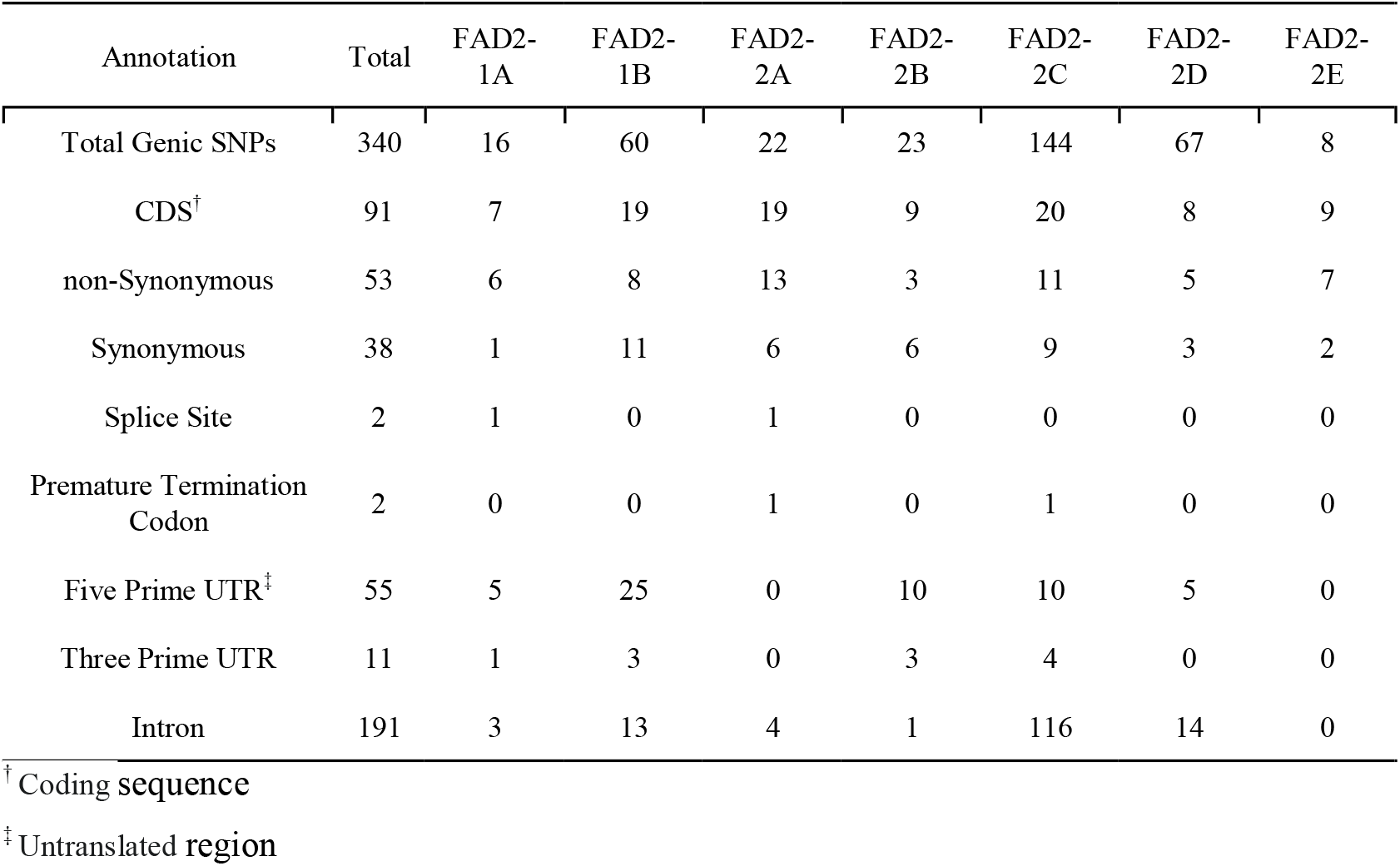
Total SNPs identified in *FAD2* family genes using the 32mSNPs

| Annotation | Total | FAD2-1A | FAD2-1B | FAD2-2A | FAD2-2B | FAD2-2C | FAD2-2D | FAD2-2E |
| --- | --- | --- | --- | --- | --- | --- | --- | --- |
| Total Genic SNPs | 340 | 16 | 60 | 22 | 23 | 144 | 67 | 8 |
| CDS<sup>†</sup> | 91 | 7 | 19 | 19 | 9 | 20 | 8 | 9 |
| non-Synonymous | 53 | 6 | 8 | 13 | 3 | 11 | 5 | 7 |
| Synonymous | 38 | 1 | 11 | 6 | 6 | 9 | 3 | 2 |
| Splice Site | 2 | 1 | 0 | 1 | 0 | 0 | 0 | 0 |
| Premature Termination Codon | 2 | 0 | 0 | 1 | 0 | 1 | 0 | 0 |
| Five Prime UTR<sup>‡</sup> | 55 | 5 | 25 | 0 | 10 | 10 | 5 | 0 |
| Three Prime UTR | 11 | 1 | 3 | 0 | 3 | 4 | 0 | 0 |
| Intron | 191 | 3 | 13 | 4 | 1 | 116 | 14 | 0 |

<sup>†</sup> Coding sequence
<sup>‡</sup> Untranslated region

**Table 3.** Accessions carrying the DNA variant (reference allele C to alternative allele G) in *FAD2-1B* that results in amino acid change at P137R.

| PI | Common Name | Species | MG | P137R |
| --- | --- | --- | --- | --- |
| PI518671 | Williams82 | *G. max* | III | C |
| PI283327 | Pingtung Pearl | *G. max* | V | G |
| PI506933 | Kouiku 1 | *G. max* | IV | G |

## Discussion

It is a critical step to make a large quantity of genome data publicly available in a user-friendly format for post-genomic studies in the era of genetic diversity (Kersey 2019). It has been long acknowledged that soybean population contains a vast amount of genetic diversity (Sun *et al*. 2015). However, analyzing a large quantity of combined genome sequences requires the investment of tremendous time and effort, particularly proficient bioinformatics skills and high-performance computing. In the study, we integrated and analyzed a total of 208.3 billion genome sequencing reads of the 1,556 accessions and identified and comprehensively annotated 32 million of SNPs present in the 1.5K soybean accessions. The genotyping data at 32mSNP positions for the 1500 diverse soybean accessions at an average density of 30 SNPs/Kb genome sequence and 12 non-synonymous SNPs/gene enable to conduct a broad range of genomic analyses at a single nucleotide resolution to advance soybean research. We also characterized the population structure and demonstrated that the collection of 1.5K accessions can be effectively used to map trait QTL or gene candidates, and identify known and unknown alleles for the trait genes in 1.5K accessions through the fine GWAS of two *Dt1* regions and *in-silico* genotyping *FAD2* gene family. Identification of genomically “equivalent”/closely related accessions between 1.5K sequenced accessions and 20K US collections of soybean accessions offers an opportunity to maximize the uses of the genomic data, allowing to access useful variation existing in the oversea accessions that are inaccessible to each other in the context of policy barriers for international germplasm exchange. Despite the dramatic drop of SNP number after MAF-based filtration, we demonstrated that retaining rare alleles in the collection is useful to identify rare but valuable alleles or accessions, as exampled by the identification of two P137R-carrying accessions (MAF = 0.12%). In addition, usage of the SNP collection may beyond what we presented here depending on research aims. For example, the 1.5K covers majority of diversity in 20K collection, imputating the Soy50KSNP data of the 20K accessions with the 32mSNPs may enable in-depth estimation of the entire USDA Soybean Collection (Arouisse *et al*., 2020). To make full use of this versatile SNP collection, we made it and the detailed annotation information public available at SoyBase (https://soybase.org) and Ag Data Commons (https://doi.org/10.15482/USDA.ADC/1519167) for convenient uses.

Enriching genetic diversity by integrating diverse gene pools is critical to discover and use novel alleles for continuous crop improvement, and this becomes increasingly important for soybean because cultivated soybean has narrow genetic basis due to severely selection during the domestication and modern breeding processes (Arsenault-Labrecque *et al*., 2018; Cheng *et al*., 2017; Patil *et al*., 2019). There are increasing evidence in various studies showing that crop wild relatives have a broader repertoire of genes/alleles that can serve as a valuable source of genetic diversity for crop improvement (Zhang *et al*., 2018), especially those lost in cultivated soybean (Maldonado dos Santos *et al*., 2016; Valliyodan *et al*. 2016; Zhou *et al*. 2015). Our study for the first time combined all the genome data of worldwide-grown accessions and integrated the representative set of 91 diverse *G. soja* accessions into the analysis, thus generating a highly dense SNP map consisting of 32 million SNPs at 1 SNP per 30 bp with an average of 12 non-synonymous SNPs/gene. In addition, the collection contains soybean accessions important to soybean improvement, such as Lee (PI548656) and Essex (PI548667) and Harosoy (PI548573) that have been served as important breeding materials in Southern and Northern US breeding programs respectively, so valuable variants important to many agronomically-important traits were included in this collection. Therefore, this 32mSNPs may serve as a core and representative collection that should be effective in advancing forward and reverse genetic studies at a high resolution of both gene and nucleotide levels. Indeed, we identified rare-occurred nucleotide substitutions in *FAD2* genes critical in altering fatty acid profiles in multiple accessions (**Tables 2** and **3**); *Dt1* was successfully fine-mapped on chr19 within a high confidence interval. We were also able to identify another QTL on chr03 that co-located with *Dt1* ortholog (**Figure 5**). Thus, the 32 million SNPs in the 1,556 diverse soybean accessions could serve as a highly versatile resource for a broad range of genomics-driven gene discovery studies, and hence providing a rich source to mine soybean diversity for valuable trait genes/variants. The collection of 1.5K diverse and representative genomes is also serving as a core to include more genome sequences while available and more annotation such as structural variations in addition to SNPs. Eventually, we can advance it to a comprehensive, large, and user-friendly genomic resource for the research community.

## Materials and methods

### Genome sequencing data collection and generation

Genome sequencing reads of all available *G. soja* and *G. max* accessions were retrieved from the Short Read Archive (SRA) database at NCBI (www.ncbi.nlm.nih.gov). The sequencing reads were generated using Illumina technology platforms. The downloaded short-read archive format was converted to zipped *fastq* format using the fastq-dump of sratoolkit. A total of 91 representative *G. soja* were sequenced in 150-bp paired-end reads using the Illumina HiSeq2000 sequencer and integrated with the downloaded sequencing reads for further analysis. Based on germplasm information, we identified that 56 accessions were sequenced twice in two different studies and one was sequenced three times by different laboratories. Considering the possibly genetic variation in the accessions with the same ID from different labs, we retained the duplicated accessions in our analysis and counted as individual accessions. The information on accession ID or name, species, maturity group, origin, and geographical origin was retrieved from the Germplasm Resources Information Network (GRIN, https://www.ars-grin.gov/) and listed in **Table S1**.

### SNP calling and annotation

The raw sequencing reads were aligned on the soybean reference genome (*G. max* cv. Williams 82.a2 v1) from Phytozome v10 (Goodstein *et al*., 2012; Schmutz *et al*. 2010) using burrows-wheeler aligner (BWA) (version: 0.7.17-r1188) (Li and Durbin 2009). Picard tool was used to add reads group, reorder and sort reads, and mark duplicated reads (version: 2.9.2, https://broadinstitute.github.io/picard/). SNPs were called using UnifiedGenotyper function in Genome Analysis Toolkit (GATK) (Version: 3.4) with default parameters (DePristo *et al*., 2011). SNPs in each sample were filtered as follows: coverage > 20%, read depth ≥ 5 reads, SNPs quality ≥ 50, and ≤ 2 SNPs in a 10-bp window. Each sample was then genotyped individually at each of the SNP sites passing the filtering criteria. SNP sites for the 1.5K collection were merged and further filtered on minor allele frequency > 0.001 and missing rate < 0.5. The genome-wide distribution of the SNPs was illustrated using Circos (Krzywinski *et al*., 2009). A custom Perl script was developed to annotate SNPs based on the coordinates of features (intron, exon, UTR) of all gene models in the reference genome (Goodstein *et al*. 2012; Schmutz *et al*. 2010). The effects of non-synonymous SNPs on gene functions were predicted by Protein Variation Effect Analyzer (PROVEAN version: 1.1.5) (Choi *et al*. 2012).

### SNP data comparison

Genotyping results by SoySNP50K iSelect Bead Chip for 20,087 (20K) soybean accessions in the US Soybean Collection were downloaded from SoyBase (https://soybase.org) (Song *et al*., 2013). Using custom Perl scripts, the SNP coordinates shared by both SoySNP50K iSelect Bead Chip and our 32mSNPs were identified and used to compare genotyping data from two technology platforms. Coordinates where either no call or N at the shared coordinates were not included in the comparison. The percent of identity in the comparison was calculated using the formula:

Percent of Identity = (Sum of Identical SNPs/All Coordinates Compared Count) * 100%

The 15,623,492 registered soybean SNP coordinates in The NCBI’s dbSNP Database (https://www.ncbi.nlm.nih.gov/SNP) were downloaded and compared with the 32mSNPs of 1.5K genomes. The coordinates in 32mSNPs not matching with those in dbSNP database were regarded as newly identified SNPs.

### Population genomics analysis

Principal component analysis (PCA) was used to analyze the population structure and relation between re-sequenced 1,556 accessions and the 20,087 accessions in the GRIN database using filtered 32mSNPs and SoySNP50K SNPs, respectively. PCA was conducted using the SNPRelate package (Zheng *et al*., 2012). An identity-by-state (IBS)-based neighbor-joining (NJ) phylogenetic tree was constructed for all accessions using an R package SNPRelate (Zheng *et al*. 2012). The phylogenetic tree was visualized using *ape* packages (Paradis et al., 2004). All SNPs used in this analysis were filtered using *snpdsLDpruning* function in SNPRelate with ld.threshold = 0.5, missing.rate = 0.2. LD decay was analyzed using PopLDdecay with -MaxDist = 1000 and -MAF = 0.05 (Zhang *et al*., 2019).

### Genome-wide association study

The phenotypic data for stem determinate was downloaded from the GRIN database. SNP data were filtered to keep only biallelic SNP and minor allele frequency greater than 0.01. GWAS was performed using a linear mixed model in GAPIT (Lipka *et al*., 2012; Yu *et al*., 2006) with Model.selection =T. Kinship was calculated using the default setting in GAPIT and included in the analysis. The threshold for significant association was determined with *p* < 0.05/total amount of SNPs used in the analysis. The regional LD heatmap was generated using the LDheatmap package (Shin *et al*., 2006).

## Acknowledgment

The authors would like to acknowledge Rick Meyer for his critical technical support in computationally data processing and analysis, Dr. Gunvant Patil for helps in drafting the manuscript, Drs. Anne Brown, Steven Cannon, Nelson Rex to release the data on SoyBase, and many researchers for deposing the genome sequencing data in the public to make the research possible, and Dr. Dilip Shah for his kind help with this project. The research is funded by the United Soybean Board (USB #2020-162-0202), USDA-ARS (Project #: 5070-21000-042-00-D) and Bayer AG (Appl. No: 2019-01-058).

## Author contribution

YQA concepted the study, participated in data analysis and drafted the manuscript. HZ, HJ and ZH drafted manuscript and participated in data analysis; and QS generated genome sequencing and participated in drafting manuscript.

## Disclaimer note

Names are necessary to report factually on available data; however, the USDA neither guarantees nor warrants the standard of the product, and the use of the name by USDA implies no approval of the product to the exclusion of others that may also be suitable. USDA is an equal opportunity provider and employer.

## Table Legends

**Table S1** Information of accession ID, origin, and statistical result for the 1.5K collection

**Table S2** Best match of each of the 20K accessions to the 1.5K accessions

